# *Sonic hedgehog* is essential for proximal-distal outgrowth of the limb bud in salamanders

**DOI:** 10.1101/2021.09.16.460693

**Authors:** Sruthi Purushothaman, Brianda B. Lopez Aviña, Ashley W. Seifert

## Abstract

The developing forelimb has been a foundational model to understand how specified progenitor cells integrate genetic information to produce the tetrapod limb bauplan (*1, 2*). Although the reigning hypothesis is that all tetrapods develop limbs in a similar manner, recent work suggests that urodeles have evolved a derived mode of limb development (*3-5*). Here we demonstrate through pharmacological and genetic inactivation of *Sonic hedgehog* (*Shh*) signaling in axolotls that *Shh* directs expansion and survival of limb progenitor cells in addition to patterning the limb across the proximodistal and antero-posterior axis. In contrast to inactivation of *Shh* in mouse or chick embryos where a humerus, radius and single digit develop (*6-9*), *Shh* crispant axolotls completely lack forelimbs. In rescuing limb development by implanting SHH-N protein beads into the nascent limb field of *Shh*-crispants, we show that the limb field is specified in the absence of *Shh* and that hedgehog pathway activation is required to initiate proximodistal outgrowth. When the derived nature of salamander limb development is placed in a phylogenetic context, it generates a new hypothesis where the ability to regenerate an entire tetrapod limb may have evolved uniquely among urodeles.

**Teaser:** *Shh* is essential for salamander limb development

## Introduction

Genetic and molecular investigation of early amniote forelimb and pectoral fin development has revealed a high degree of mechanistic conservation across the relatively few model organisms that have been well studied (e.g., mouse, chick, *Xenopus*, zebrafish). Forelimb development in amniote embryos, and to a large extent pectoral fin development, can be deconstructed into four general phases: progenitor field establishment and positioning, initiation and expansion of limb progenitor cells, patterning of limb progenitors across three cardinal axes (anteroposterior, proximodistal, dorsoventral) and morphogenesis of the mesoderm into a mature limb integrating muscles, skeletal elements and connective tissues (*2, 10, 11*). Forelimb and pectoral fin field establishment begins when retinoic acid (RA) specifies a subpopulation of the somatopleure (*12-15*) to become forelimb/fin mesoderm while *Hox* gene expression aligns these progenitors along the craniocaudal (head-to-tail) axis (*16, 17*). Forelimb and pectoral fin bud initiation occurs when RA and canonical Wnt-signaling subsequently induce *Tbx5* among forelimb/pectoral fin field progenitors (*13, 18-22*). Limb and fin bud outgrowth occurs when *Fgf10* is activated throughout the nascent bud mesoderm which induces Fgf-signaling in the overlying ectoderm to create positive feedback between the ectoderm and mesoderm (*23-27*). As the forelimb and pectoral fin bud emerges from the body wall it acquires anteroposterior polarity with *Hand2* and several *HoxA/D* genes restricted to the posterior and *Gli3* expression portioned into the anterior mesoderm (*28-32*). Subsequently, two signaling centers form that control limb development along the proximodistal and anteroposterior axes respectively: the apical ectodermal ridge (AER), marked primarily by *Fgf8* expression, and the zone of polarizing activity (ZPA), marked by *Sonic hedgehog* (*Shh*) expression (*33-38*).

AER excisions in chicken embryos and gene knockout experiments in mice demonstrated that fibroblast growth factors (Fgfs) secreted from the AER are absolutely essential for limb development where they promote cell survival and proximodistal outgrowth of the limb (*33, 39-41*). Inactivation of *Fgfs 4, 8* and *9* in the ectoderm (*39*) or early removal of limb bud ectoderm results in a scapula alone (*33*). Similarly, the role of the ZPA and Shh-signaling has been extensively studied during chicken and mouse limb development. Following outgrowth and establishment of the AER, Shh sets up anteroposterior positional values (*35, 42, 43*), maintains AER width and expression of the AER-*Fgfs* (via *Gremlin1* restriction of Bmp-signaling) (*8, 9, 44-47*), and regulates cell proliferation (*48-50*) and cell survival of limb mesoderm (*51*). The spatial restriction of AER-*Fgfs* and *Shh* have been analyzed in a spectrum of vertebrate species supporting conserved expression in the ectoderm and mesoderm respectively (*35-38, 52-54*).

Despite this apparent conservation, previous work has shown that salamanders lack an AER (*4, 5*) and that at some point during amphibian evolution localization of *Fgfs* and *Fgf* receptors shifted to the limb mesenchyme where they now control limb size but are largely dispensable for limb development (*3*). At the very least, these data suggest that the molecular logic of tetrapod limb development may not be absolutely conserved among all tetrapods. Here we asked whether reduced prominence of Fgf-signaling during salamander limb development might be offset by an increased reliance on Shh-signaling to control limb field progenitor proliferation, survival, outgrowth and patterning. To test our hypothesis, we pharmacologically inhibited Shh-signaling in axolotls using cyclopamine or the highly specific smoothened antagonist BMS-833923. We also genetically inactivated *Shh* using CRISPR/Cas9 and analyzed limb development in *Shh* crispants. We asked whether SHH-N or FGF8 protein could stimulate limb development outside the limb field as has been observed in chick embryos (*55*) and whether the limb field formed in the absence of Shh signaling.

## Results

### Small molecule smoothened antagonist BMS-833923 inhibits proximodistal outgrowth of the limb bud in axolotl embryos

To test the hypothesis that Shh-signaling coordinates expansion of limb field progenitor cells and proximodistal outgrowth of the limb bud in salamanders, we first inhibited Shh-signaling throughout early limb development using a pharmacologic approach. Previous studies using cyclopamine during axolotl limb development provided evidence that *Shh* functions primarily to pattern the anteroposterior axis following expansion of limb bud progenitors (*56*). Curiously, recent work exploring Shh-signaling during zebrafish development and caudal fin regeneration revealed that a highly specific smoothened antagonist, BMS-833923 (hereafter BMS) (*57*) more potently and specifically inhibited hedgehog signaling compared to cyclopamine (*58*). Therefore, we treated pre-limb bud stage larvae (stage 39) with ethanol, cyclopamine or BMS for ten days (initiation and expansion phase) and harvested limbs at stages 46 and 54 (Figure 1A-B and Figure S1A-C). Analyzing skeletal differentiation at stage 54, we found cyclopamine treatment primarily affected anteroposterior patterning, with nearly all (∼88%) resultant limbs possessing a humerus, single fused radius/ulna and at least one digit (Figure 1B and Figure S1A, B & D). These results mirrored previous limb development studies in urodele and amniote embryos using the same concentration of cyclopamine (*9, 56, 59*). In stark contrast, 92% of the BMS-treated larvae had no limbs with only a small bump covering the scapula where the humerus would normally articulate or no bump at all (Figure. 1B and Figure. S1C, D).

**Figure 1.**
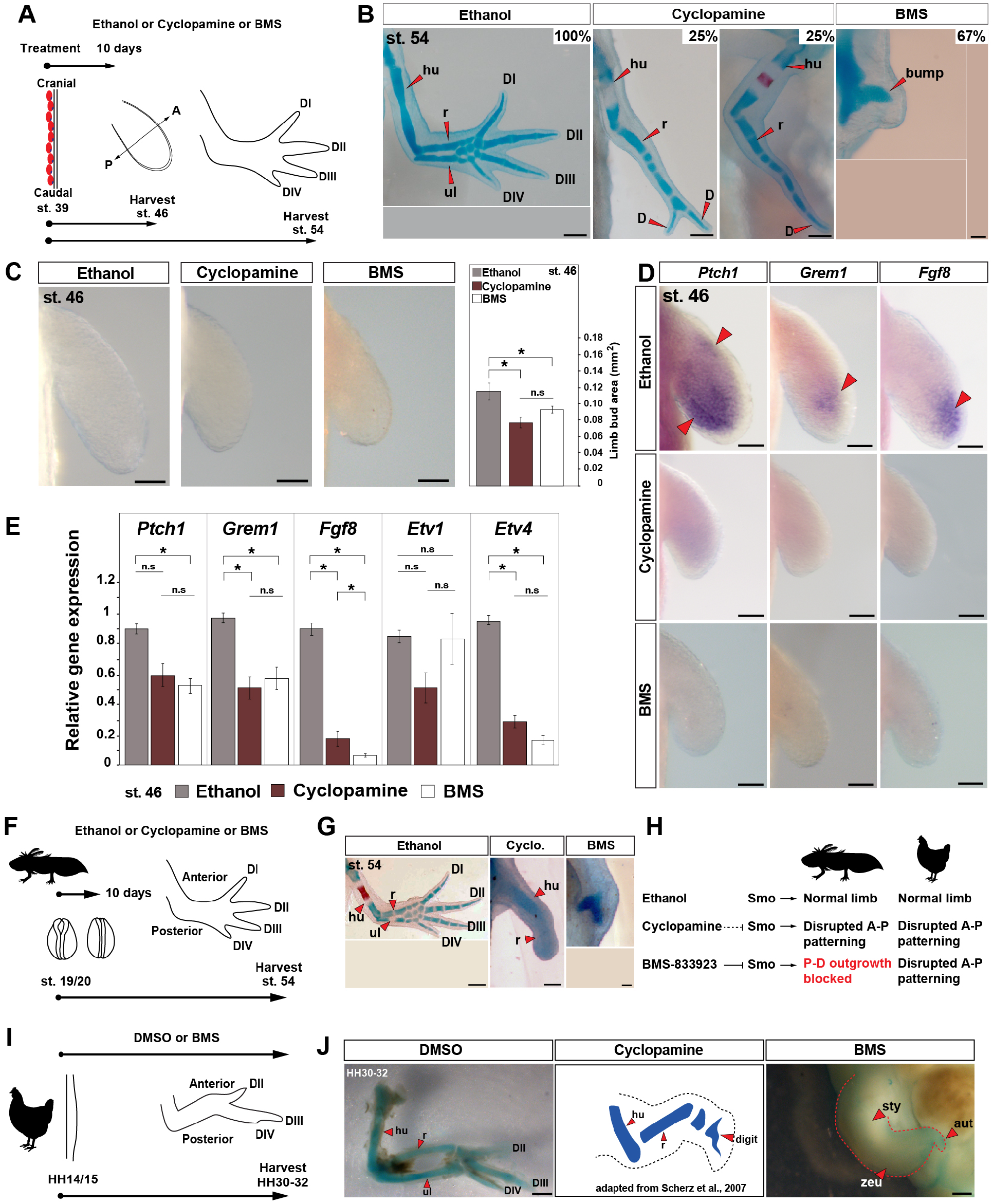
Small molecule smoothened antagonist BMS-833923 inhibits limb bud outgrowth in axolotl larvae. **(A)** Design for ethanol (control), cyclopamine and BMS treatments in axolotl. Limbs are aligned with anterior ‘A’ on the top and posterior ‘P’ on the bottom. Red ovals depict dorsal muscle blocks. (**B**) Representative images of Alcian blue/Alizarin red stained ethanol, cyclopamine or BMS treated stage 54 limbs (limb n=30 for ethanol, =24 for cyclopamine and =36 for BMS-833923). Scale bar = 500 μm. (**C**) Limb bud area measurements in ethanol (control), cyclopamine or BMS treated limbs (n=6/treatment). Scale bar = 100 μm. (**D**) *In-situ* hybridization for genes *Ptch1, Grem1* and *Fgf8* in ethanol, cyclopamine or BMS treated stage 46 limbs (n=3 or 4/gene). Red arrows: expression domain. Scale bar = 100 μm. (**E**) qRT-PCR of the *Ptch1, Grem1, Fgf8, Etv1* and *Etv4* expression in stage 46 limbs post ethanol, cyclopamine or BMS treatments (n=3/treatment). (**F)** Design for ethanol, cyclopamine or BMS treatments at neural fold stage 19/20 in axolotls. **(G)** Alcian blue/Alizarin red staining at stage 54 for neural fold treatments with ethanol, cyclopamine or BMS (n=3/ treatment). Scale bar = 500 μm. **(H)** Schematic depicting the mode of actions of ethanol, cyclopamine and BMS in axolotl and chick limb buds. **(I)** Design for ethanol, cyclopamine or BMS treatments at HH14/15 in chick embryos. **(J)** Alcian blue/Alizarin red stained DMSO, cyclopamine (adapted from Scherz et al., 2007) and BMS treated limbs at HH30-32 (n=4/treatment). Scale bar = 1mm. Error bars: SEM and asterisk: significant P-values. hu, humerus; r, radius; ul, ulna; D, digit; sty, stylopod; zeu, zeugopod and aut, autopod.

Examining pre-chondrogenic limbs at stage 46, we observed that treatment with cyclopamine or BMS caused a significant decrease in limb bud size, although small limb buds still formed in both treatment groups (One-way ANOVA, Tukey-Kramer HSD post hoc test, *F*=8.628; Tukey-Kramer HSD post hoc test, *P*=0.0026 (ethanol vs. cyclopamine), *P*=0.046 (ethanol vs. BMS) and *P*=0.34 (cyclopamine vs. BMS), n=6/treatment) (Figure 1C). To assess the degree to which these drugs inhibited Shh-signaling, we assessed expression of the direct target gene *Patched1* (*Ptch1*) and downstream target genes, *Gremlin1* (*Grem1*) and *Fibroblast growth factor 8* (*Fgf8*) in stage 46 limb buds (Figure 1D, E). Using *in-situ* hybridization and qRT-PCR, we observed that *Ptch1* was significantly downregulated by BMS treatment, *Grem1* was significantly downregulated in BMS and cyclopamine treated limbs and *Fgf8* was significantly downregulated in BMS compared to cyclopamine treated limbs (One-way ANOVA, Tukey-Kramer HSD post hoc test; *Ptch1*: *F*=7.98, *P*=0.06 (ethanol vs. cyclopamine), *P*=0.02 (ethanol vs. BMS) and *P*=0.65 (cyclopamine vs. BMS); *Grem1*: *F*=8.65, *P*=0.018 (ethanol vs. cyclopamine), *P*=0.048 (ethanol vs. BMS) and *P*=0.68 (cyclopamine vs. BMS); *Fgf8*: *F*=301.43, *P* <0.0001 (ethanol vs. cyclopamine), *P*<0.0001 (ethanol vs. BMS) and *P*=0.03 (cyclopamine vs. BMS); n=3/treatment) (Figure 1D, E). qRT-PCR for downstream targets of Fgf-signaling, ETS transcription factor family genes *Etv1* and *Etv4* showed that *Etv4* was significantly downregulated in both the drug treatments (Figure 1E) (ANOVA, Tukey-Kramer HSD post hoc test; *Etv1*: *F*=2.986, *P*=0.16 (ethanol vs. cyclopamine), *P*=0.99 (ethanol vs. BMS) and *P*=0.17 (cyclopamine vs. BMS); *Etv4*: *F*=225.92, *P*<0.0001 (ethanol vs. cyclopamine), *P*<0.0001 (ethanol vs. BMS) and *P*=0.13 (cyclopamine vs. BMS); n=3/treatment). These data supported that BMS was more effective at inhibiting Shh-signaling compared to a max dose of cyclopamine. Although BMS drug treatment ultimately resulted in no proximodistal outgrowth, analysis of initial limb bud size and expression of direct downstream targets at stage 46 revealed that neither drug completely inhibited Shh-signaling when treated at stage 39.

Although we used a max dosage of cyclopamine that embryos could tolerate without lethality (twice the concentrations used in previous amphibian studies), to rule out any potential for delayed activity of cyclopamine (compared to BMS) we exposed embryos to the two drugs prior to limb field formation (Figure 1F). Embryos treated at neural fold stage 19/20 with cyclopamine still developed a humerus and radius while BMS completely inhibited limb formation (Figure 1F-H). These data demonstrate that BMS more completely inhibits the Shh-signaling pathway compared to cyclopamine when used on salamander embryos and that Shh-signaling regulates the earliest stages of axolotl limb development similar to phenotypes recovered from Shh inactivation during pectoral fin development in zebrafish (*54*). In addition, the incomplete inhibition we observed using cyclopamine allowed us to confirm that Shh also governs anteroposterior patterning across the limb mesoderm after bud outgrowth (*49*).

Differences across the drug treatments raised the possibility that previous cyclopamine studies may have overlooked an early role for Shh-signaling in chick limbs. To test this idea, we inhibited Shh-signaling during chick limb development using BMS beginning at stage HH14 to exclude the possibility of hedgehog pathway activation prior to the treatments (Figure 1H-J). In contrast to our results in axolotls, chicken embryos treated with BMS developed limbs with a stunted stylopod, zeugopod and autopod similar to the most severely affected pair of wings in a previous study using cyclopamine (*9*) (Figure 1J). These findings substantiate that Shh-signaling in amniote limbs functions primarily to pattern the anteroposterior axis where it acts subordinately to the AER which controls proximodistal outgrowth and skeletal differentiation (Figure 1H). Our findings in axolotls, however, show that Shh-signaling first coordinates expansion of limb progenitor cells and proximodistal outgrowth of the limb bud in salamanders (Figure 1H).

### Axolotl *Shh* crispants completely lack forelimbs

To further interrogate the function of Shh-signaling during salamander limb development we genetically inactivated *Shh* in fertilized axolotl zygotes using CRISPR/Cas9. By designing three complementary guide RNAs to the *Shh* locus and separately injecting these into fertilized single cells, we recovered overlapping and robust mutant phenotypes using all three guide RNAs thereby ruling out the chances of off-target effects (Figure 2A-B and Figure S2A, Figure S3). Next generation sequencing (NGS) confirmed the efficiency of the three guide RNAs to create a frame mutation rate of ∼99% allowing us to analyze F0 larvae (Figure S2B, Table S2). Out of 300 embryos injected with either *Shh*-guide RNA#1, 2 or 3, 67 survived and >80% of the F0 crispants we screened presented a range of severe craniofacial defects, including partial to complete cyclopia, caudal truncations, and a curved body axis (Figure 2A-B); phenotypes that were similar to those observed in *Shh* null mice (*6, 7*). Although complete cyclopia occurred in relatively few F0 crispants (3%), 87% of the crispants exhibited eyes that were positioned with little to no interocular separation (Figure 2A-B). This resulted in a tight correlation between reduced interocular distance and the appearance of a bulge on the front of the head similar to the formation of a proboscis in *Shh* null mice (*6*). *Shh* mutants also had smaller heads and lacked most of the anterior craniofacial skeleton including jaws (Figure 2A).

**Figure 2.**
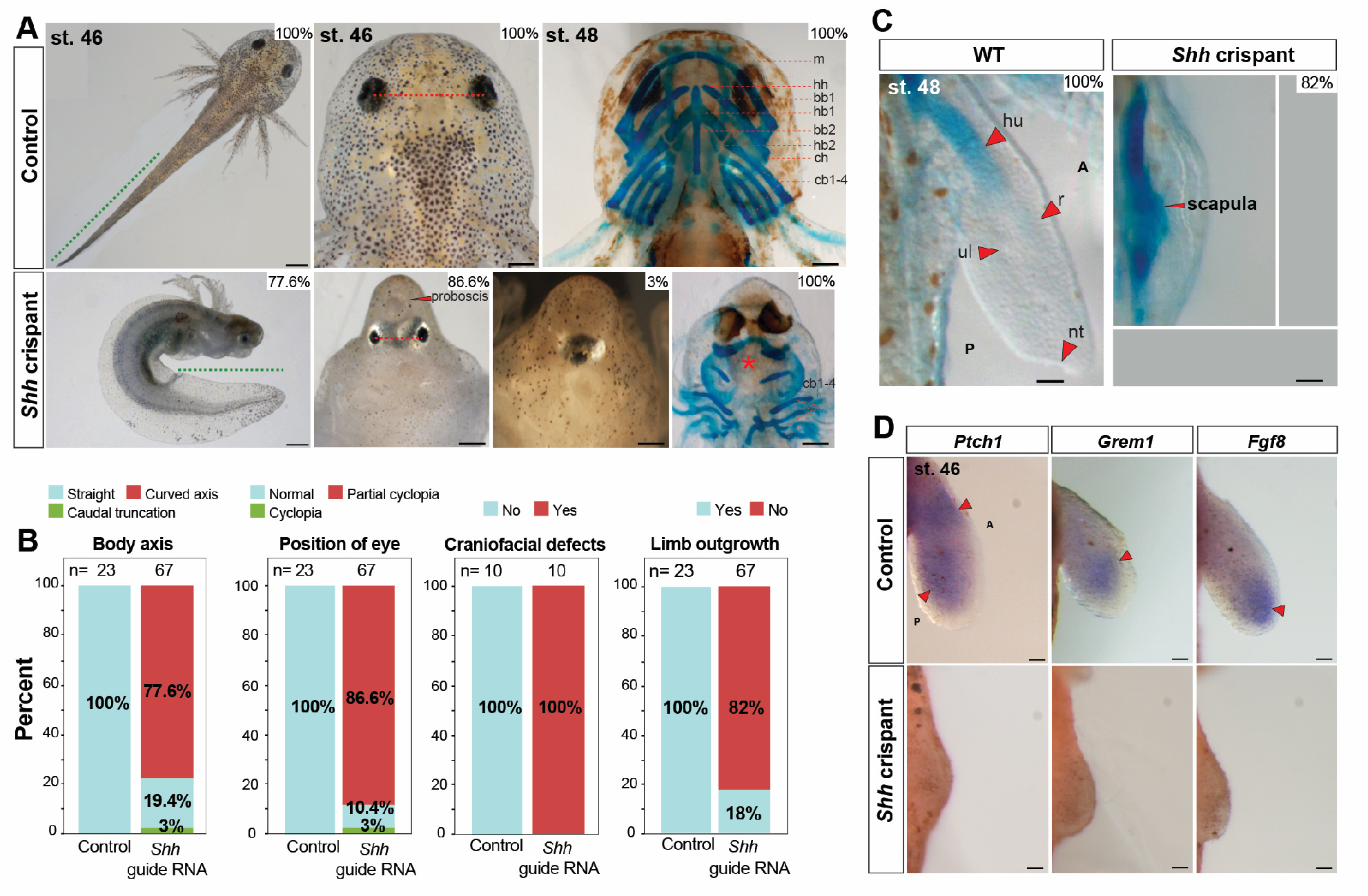
Axolotl *Shh* crispants completely lack forelimbs. (**A, B**) Phenotypes like body axis, position of the eye and Alcian blue/Alizarin red staining of craniofacial structures of CRISPR control and *Shh* crispant larvae (n=23 for CRISPR control and n=67 for *Shh* crispant larvae for body axis and eye position analysis, n=10 each for CRISPR control and *Shh* crispant larvae for Alcian blue/Alizarin red staining of craniofacial structures). Green dotted line: tail length, red dotted line: distance between the eyes, red asterisk: loss of anterior cranio-facial structures. Scale bar = 1 mm (for body axis), = 500 μm (for eye position and cranio-facial structures). (**B, C**) Alcian blue/Alizarin red staining for stage 48 limbs of CRISPR control and *Shh* crispant larvae (n=10). Scale bar= 500 μm. (**D**) *In-situ* hybridization for genes *Ptch1, Grem1* and *Fgf8* in CRISPR control and *Shh* crispant larvae at stage 46 limbs (n=3 or 4/treatments). Red arrows: expression domains. Scale bar = 100 μm.

In addition to these defects, almost all *Shh* crispants completely lacked forelimbs, a phenotype similar to zebrafish *Shh* mutants, but in contrast to *Shh* null mice, and chick embryos treated with BMS (Figure 1J and 2B-C) (*6, 7, 54*). Owing to a lack of jaw structures in the knockout animals which precluded them from eating, we analyzed the limb skeletons at stage 48 just prior to the onset of feeding (Figure 2C). At stage 48, 100% of control larvae showed a chondrifying humerus, radius and ulna while 82% of the *Shh* knockouts showed only elements of the pectoral girdle (Figure 2C). Using a previously published guide RNA against *Tyrosinase* (*Tyr*) (*60*), we observed loss of pigmentation in all injected animals, but otherwise normal embryos; a result which reinforced the specificity of CRISPR/Cas9 in axolotl embryos (Figure S4A-E). Next, we analyzed the limb field area in *Shh* knockout animals at stage 46 where limb buds should form and observed almost no outgrowth over the condensing pectoral skeleton compared to elongate limb buds observed in wildtype animals (Figure 2C, D). This outgrowth defect was even more pronounced than in our BMS treatments with ectoderm almost directly covering the scapula. Lastly, we analyzed downstream Shh targets in the limb forming region of *Shh* crispants at stage 46. While we observed strong expression of *Ptch1, Grem1* and *Fgf8* in control limbs, we were unable to detect expression for these target genes in the forelimb fields of *Shh* crispants (Figure 2D). Together with our BMS experiments, these results demonstrate that Shh-signaling is required to stimulate expansion of forelimb bud progenitors and control proximodistal outgrowth of the limb bud.

### Forelimb bud progenitor cells are specified in *Shh* crispants

In chicken embryos, Fgf-signaling is sufficient to induce a limb from uncommitted flank tissue supporting Fgf-signaling at the apex of a molecular limb program that can induce a secondary limb field (*55*). In order to ascertain if Shh-signaling could alone trigger limb bud outgrowth, we grafted SHH-N protein in 1X PBS + 0.1% BSA infused Affi-gel beads into the right flank of stage 37-39 axolotl embryos, several days before forelimb buds emerge from the forelimb field and monitored for development of an extra limb bud at stage 46 (Figure 3A). On the contralateral (left) side of these embryos we implanted 1X PBS + 0.1% BSA soaked beads as control for the bead implant (Figure 3A). In either case we did not observe the development of extra limb buds from the flank sites where we implanted SHH-N or control beads (Figure 3B).

**Figure 3.**
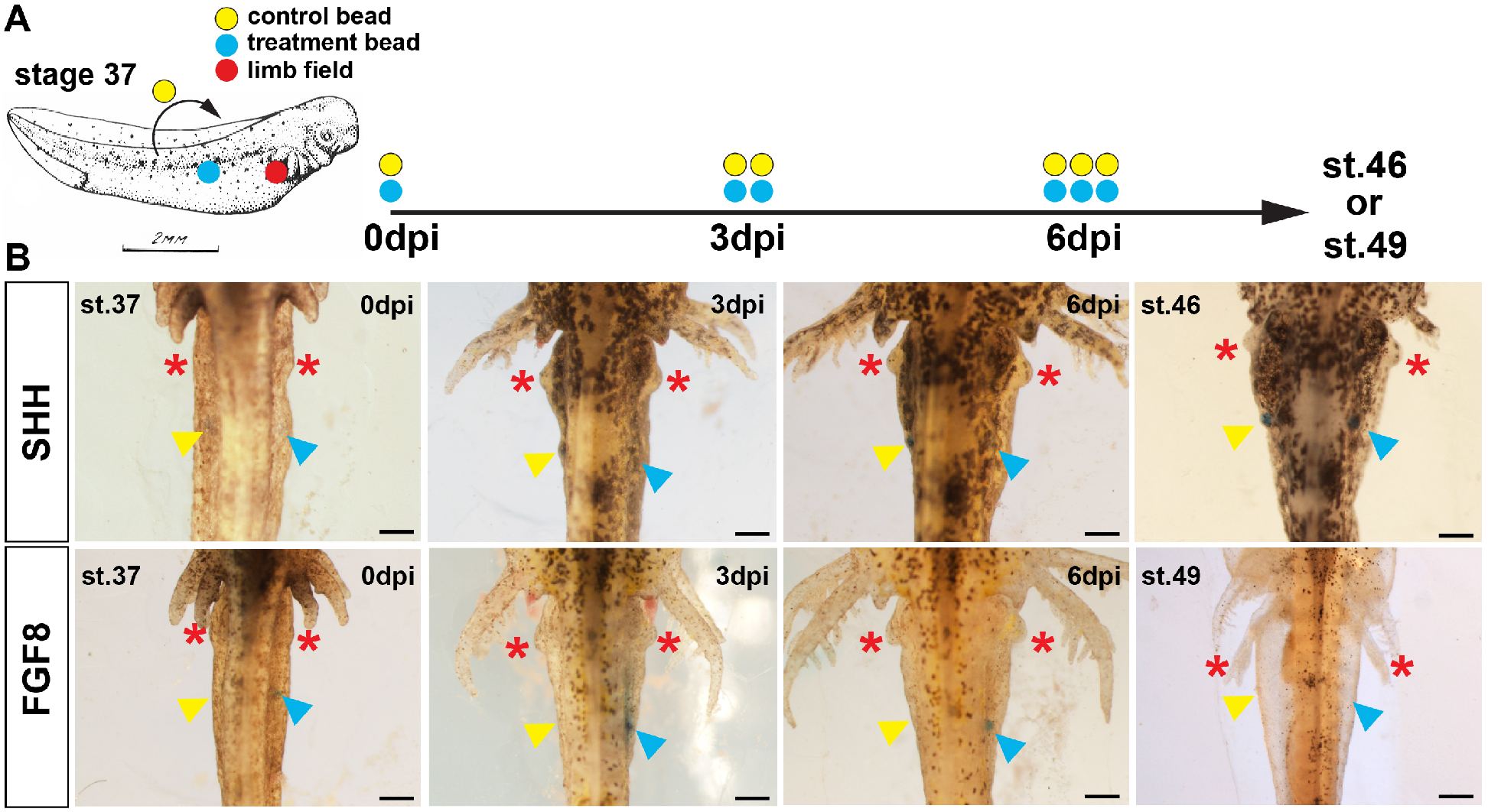
Exogenous SHH protein stimulates forelimb development from *Shh* crispant forelimb progenitors. **(A)** Schematic representation of the embryonic bead implantation experiment using stage 37 axolotl embryos. Colored circles represent the right limb field (red), relative implantation site of protein-soaked beads (blue) or PBS/0.1% BSA soaked beads (yellow). **(B-C)** SHH-N and FGF8 protein-soaked beads were grafted into the right flank lateral to the somites (blue arrow) and PBS/0.1% BSA were implanted into symmetrical positions in the left flank (yellow arrow). Three days post implantation (dpi), a second pair of beads were implanted into the same location as the first beads and consequently, a third pair were implanted at 6dpi. Evidence of ectopic limb development were tracked till stage 46 (SHH-N) or stage 49 (FGF8) and in either treatment condition we did not observe evidence of limb outgrowth. Red asterisks mark emergence and growth of the normal forelimbs. Drawing in (A) adopted from (*63*). Scale bar = 500 μm.

In contrast to the flank sites with implanted beads, normal forelimb buds were evident 3 days post implantation (dpi) (Figure 3B). This result suggested that flank mesoderm was already committed to a non-limb fate but did not rule out the possibility that another factor could induce an ectopic limb bud. Fgf8 is the endogenous inducer of chick limb formation and implanting FGF8-soaked beads can induce an ectopic limb bud and development of a complete limb (*61*). Similar to our experiment with SHH-N, we did not observe ectopic forelimb buds in response to FGF8-soaked beads and thus these data supported that limb field progenitors are specified at precise positions very early during salamander development (*62*) and that neither Shh nor Fgf8 could induce secondary limb fields in stage 37 embryos.

Based on these results, we next asked whether the forelsimb field was specified in *Shh* crispants and if so, whether implantation of SHH-N protein could induce forelimb formation from forelimb field progenitors. We implanted beads just prior to when limb buds would normally emerge i.e., stage 39 (Figure 4). Stage 39 axolotls were characterized by longer and branched gills, distinct cloaca, pigmented eyes and flanks and these features were used to roughly figure out the stage in both the control and *Shh* crispants (*63*). Affi-gel blue beads infused with SHH-N protein were implanted into the position of the forelimb field (somites 3-5) while beads containing 1X PBS + 0.1% BSA were implanted into the left contralateral forelimb field as controls and beads were replaced once every 3-4 days. While none of the beads on the control flank stimulated limb development, in 7/9 animals, limbs emerged in response to SHH-N (n=2 showed nubbin like outgrowth, n=3 showed progression to stage 45 limb bud, n=2 showed progression to stage 46/47 limb bud) (Figure 4). Of the 5/9 limbs that emerged past a nubbin and showed proximodistal outgrowth, three exhibited cartilage formation at the humerus level (Figure 4). Due to the inability of the crispants to feed, we could not take the limbs out far enough to determine if the entire limb skeleton formed. However, these data do demonstrate that forelimb progenitor cells are competent to respond to exogenously delivered SHH protein which is sufficient to stimulate expansion of forelimb progenitor cells, formation of a forelimb bud, proximodistal outgrowth of the limb bud, and skeletal differentiation of the developing limb.

**Figure 4.**
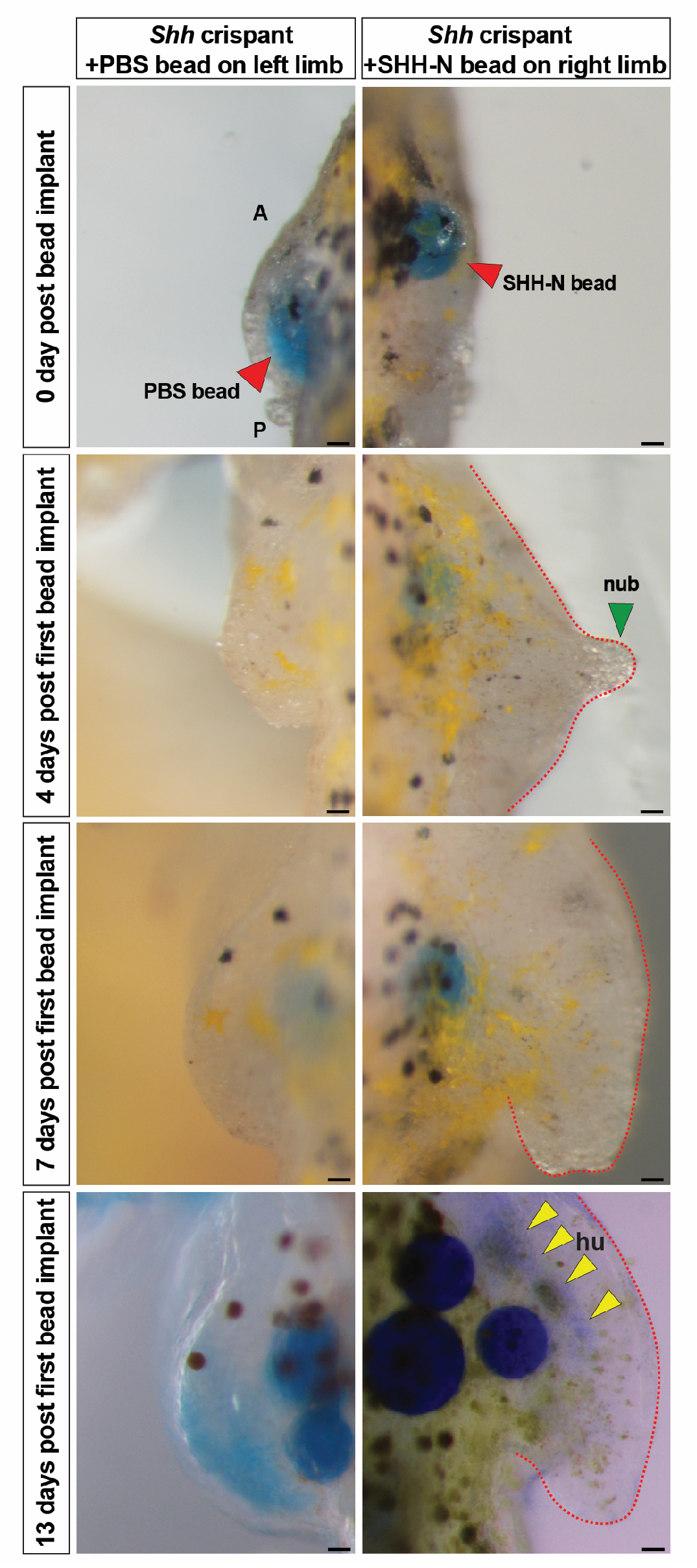
Exogenous SHH-N protein stimulates forelimb development from *Shh* crispant forelimb progenitors. Limb rescue experiments using beads loaded with SHH-N protein. Control Affi-Gel blue beads (red arrow) soaked in 1X PBS with 0.1% BSA and Affi-Gel blue beads (red arrow) soaked in 0.5 or 0.25 μg/ul SHH-N protein in 1X PBS with 0.1% BSA were grafted into the left and right limb fields respectively of *Shh* crispants. Yellow arrows: faint Alcian blue staining for humerus. Scale bar =100 μm. All the images are representative (with highest %) phenotypes. All limbs are projected in dorsal view with anterior ‘A’ on top and posterior ‘P’ on the bottom. hu, humerus and nub, nubbin.

### *Sonic hedgehog* controls cell proliferation and cell survival during axolotl limb development

Although AER-Fgf-signaling regulates cell proliferation, cell survival and limb outgrowth in chicken and mouse limbs (*39, 64, 65*), our BMS-treated and *Shh* crispant embryos suggested that Shh-signaling regulated these processes in salamanders. To address this possibility, we quantified total cell proliferation in stage 45 limb buds across all treatment groups using lightsheet microscopy (*3*). Compared to control limb buds, cyclopamine treated limbs showed a decrease in proliferating cells only at the distal tip of the limb and did not show a significant decrease in total EdU+ cells (One-way ANOVA, Tukey-Kramer HSD post hoc test, *F*=5.8, *P*=0.038 (ethanol vs. BMS), *P*=0.64 (ethanol vs. cyclopamine), *P*=0.12 (cyclopamine vs. BMS), n=3/treatment) (Figure 5A, C). In contrast, BMS-treated embryos and *Shh*-crispant embryos showed a significant decrease among total EdU+ cells compared to control limbs (Figure 5A-C) (One-way ANOVA, Tukey-Kramer HSD post hoc test, *F*=614, *P*<0.0001 (control vs *Shh*-crispant), n=3/treatment). Although the fraction of total proliferating cells as a function of total limb volume was not different between the cyclopamine and BMS treatments at stage 45, it was evident from the lightsheet images that the BMS-treated limbs were significantly smaller and contained fewer mesodermal cells (Figure 5A).

**Figure 5.**
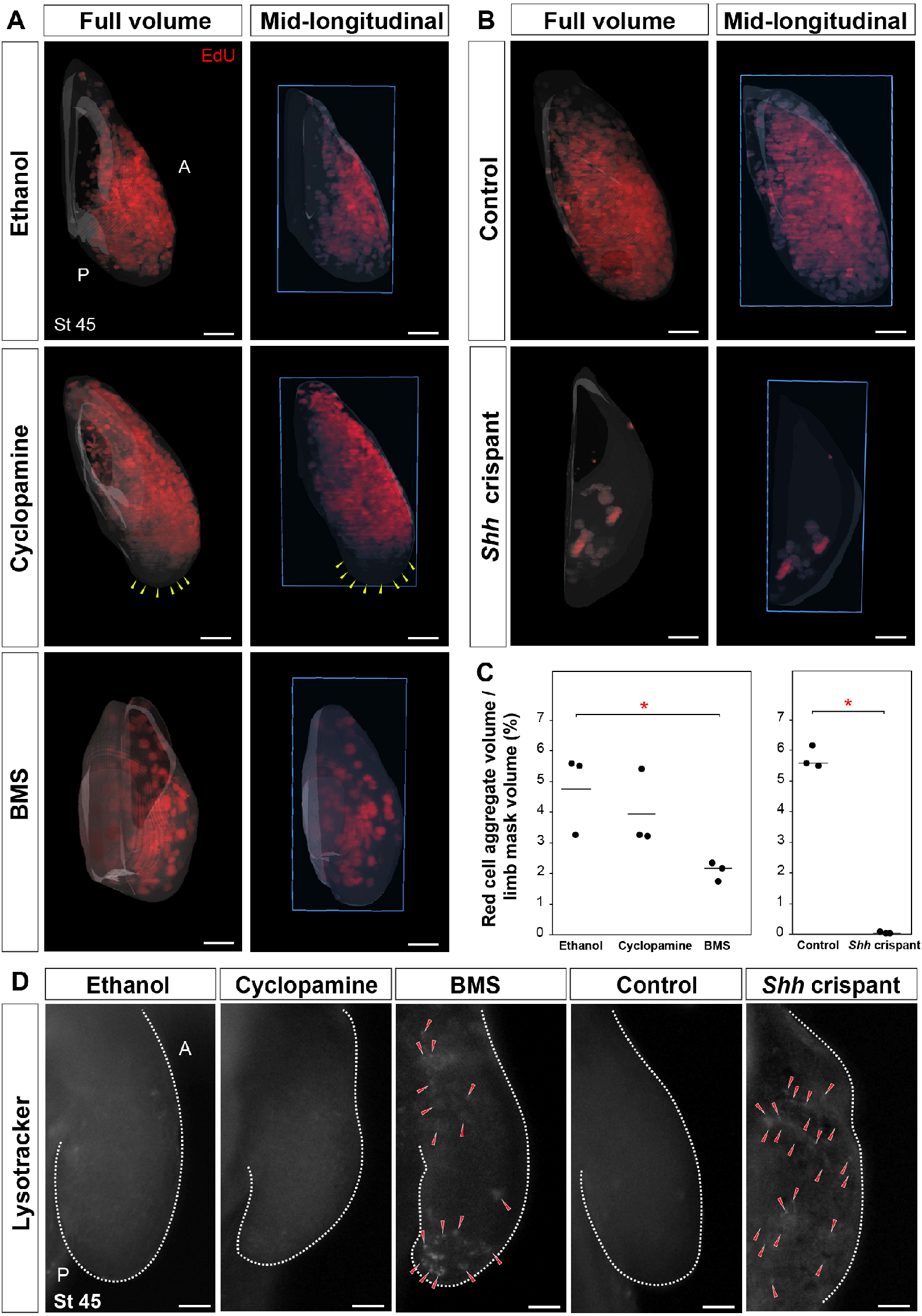
*Sonic hedgehog* controls cell proliferation and cell survival during axolotl limb development. (**A, B**) Lightsheet images depicting EdU positive proliferating cells in stage 45 limbs from ethanol, cyclopamine or BMS treatments and CRISPR control and *Shh* crispant larvae (n=3/treatment). Yellow arrows: zone lacking proliferating cells. Blue box: plane of mid-longitudinal section. **(C)** Stage 45 limbs from BMS treated larvae and *Shh* crispant larvae showed a significant decrease in EdU positive proliferating cells. Horizontal bars: mean values, asterisk: significant P-value. **(D)** Cell death assay using Lysotracker in stage 45 limb in ethanol, cyclopamine or BMS treated larvae and CRISPR control and *Shh* crispant larvae (n=3/treatment). Red arrows: lysotracker positive cells. All limbs are projected in dorsal view with anterior ‘A’ on top and posterior ‘P’ on the bottom. Scale bar =100 μm.

Next, we analyzed cell survival using Lysotracker to label dying cells within developing stage 45 limbs (*39, 66*). BMS-treated and *Shh*-crispant larvae showed Lysotracker positive cells throughout the limb buds, while none of the control larval limbs showed Lysotracker positive cells (Figure 3D). Cell death was most prominent in the proximal and distal ends of BMS-treated limbs while dying cells were present all throughout the area of the limb field present on the flanks of *Shh* crispants (Figure 5D). Together, these results from *Shh* crispants and BMS-treated axolotl larvae support a model where Shh-signaling regulates cell proliferation and cell survival in mesodermal progenitors of the limb field. This is consistent with our bead experiments in *Shh* crispants where Shh induces expansion of limb bud progenitors and is sufficient to initiate limb bud formation and outgrowth.

## Discussion

Our study supports a model where molecular components present in pectoral fin and amniote forelimb buds are deployed uniquely during salamander limb development. Specifically, our results demonstrate that Shh-signaling is essential for proliferation, survival and expansion of forelimb field progenitor cells to form a salamander forelimb bud. As the limb bud emerges from the flank, Shh-signaling stimulates *Fgf8* and downstream Fgf-signaling targets in limb mesoderm which support some cell proliferation at the distal tip of the limb bud, but that do not significantly contribute to proximodistal outgrowth of the limb (*3*). Our results also uncovered that cyclopamine does not completely inhibit Shh-signaling in salamanders when used at a maximum, non-lethal concentration, and revealed that BMS is a more effective hedgehog inhibitor in salamander embryos. Genetic inactivation of *Shh* using CRISPR/Cas9 in newly fertilized zygotes confirmed this pharmacological difference among drugs and further revealed that Shh is not required to specify forelimb progenitor cells. Together, our data demonstrate that the molecular regulation of forelimb development has independently evolved in salamanders away from a reliance on Fgf-signaling found in anuran and amniote limbs, but also found in actinopterygian pectoral fins (Fig. 6).

**Figure 6.**
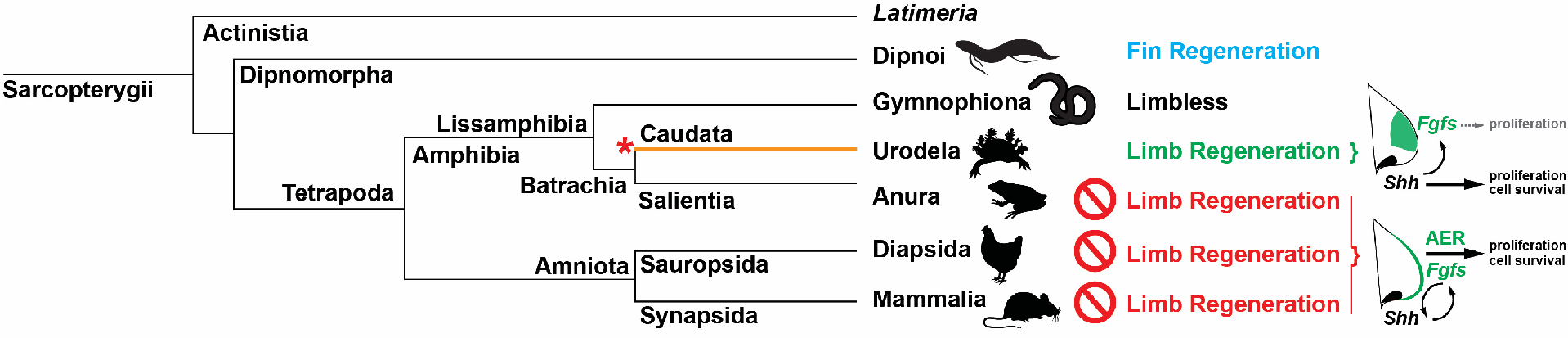
Limb regenerative ability may have evolved uniquely in caudates. Comparison of developmental mechanisms among tetrapods has revealed that urodeles possess a derived mode of limb development compared to anurans and amniotes including, absence of a morphological and molecular AER, mesenchymal restriction of canonical AER-*Fgfs*, absence of a positive feedback loop between Fgf- and Shh-signaling, and reliance on Shh-signaling to regulate proximal-distal outgrowth via cell proliferation and survival (orange line). Correlating with this unique mode of limb development, urodeles are the only extant tetrapods capable of complete limb regeneration at any stage of life (larva, juvenile or adult). The simplest explanation for this tight correlation is that limb regenerative ability evolved within caudata as their mode of limb development was modified from the ancestral tetrapod state to reside within the mesenchymal tissue compartment (red asterisk). This contradicts the hypothesis that limb and fin regenerative ability have a common origin among Sarcopterygii (*80*), a hypothesis based on common gene expression data derived from regeneration of cartilaginous elements in lungfish fins, a process similar to the regeneration of a cartilage spike in amputated *Xenopus* limbs.

Pectoral fin and forelimb development in actinopterygian and amniote embryos respectively, rely on ectodermal-mesodermal crosstalk between Fgf- and Shh-signaling. In amniote embryos where ZPA/AER crosstalk is present, proximodistal outgrowth of the limb bud is Shh-independent with Shh functioning as a cell survival factor, maintaining Fgf-signaling in the AER, and coordinating anteroposterior patterning (*8, 9, 44-47, 51*). While Shh-signaling does appear to regulate cell proliferation in chick embryos (*49*), it remains unknown if this is an indirect effect mediated through Fgf-signaling as occurs in zebrafish (*67*). Proliferation of the fin/limb mesoderm and proximodistal expansion of the bud appears reliant on Fgf-signaling from the AER which also maintains Shh-signaling from the ZPA (*39, 40, 68, 69*). For instance, although zebrafish *shh* mutants completely lack pectoral fins, fin buds emerge normally, only later to regress as ectodermal Fgf-signaling is lost (*54*). Similarly, in *Shh*^*-/-*^ mice, limb buds emerge smaller, but otherwise normally with an intact AER expressing *Fgf8* (*7*). However, without Shh-signaling from the ZPA the AER disappears, cell proliferation decreases, and posterior elements are lost (*7*). It would appear that loss of *Shh* produces a different phenotype in fishes and amniotes (i.e., no pectoral fins versus elongate forelimbs with a humerus, radius and a single digit). However, the retention of skeletal elements in amniotes stems from the extent to which Fgf-signaling is maintained in the absence of Shh-signaling to sustain mesodermal proliferation and offset cell death reinforcing the reliance on Fgf-signaling for fin and limb development in these groups. In fishes and amniotes, Fgf-signaling is required for early expansion of fin/limb bud mesoderm where genetic disruption of AER-*Fgfs* leads to a complete absence of forelimbs/pectoral fins (*25, 39, 68, 69*). Thus, Fgf-signaling from the ectoderm controls expansion of the fin/limb progenitor pool until such time that differentiation of the fin/limb skeleton begins.

In contrast to fishes and amniotes, our work demonstrates that ectodermal-mesodermal crosstalk disappeared in salamanders when Fgf-signaling moved entirely into the mesoderm (*3*); an event that also led to an erosion of the tight linkage between the two signaling pathways and an increased reliance on Shh-signaling. Importantly, one consequence of this shift was that limb development in urodeles became largely independent of Fgf-signaling. Inhibiting Fgf-signaling in salamanders produces a relatively normal limb (minus a digit) that results from an overall small decrease in cell proliferation (*3*) while inhibiting Shh-signaling in this study resulted in a complete lack of forelimbs. Although our results using BMS or genetic inactivation of *Shh* make clear that limb bud cells cannot survive without Shh-signaling, the emergence of small limb buds in response to BMS support that a miniscule amount of Shh-signaling is sufficient to stimulate cell proliferation among forelimb field progenitors, a result which could have manifested indirectly through residual Fgf-signaling (Fig. 1E).

Implantation of SHH-N loaded beads into *Shh*-crispants demonstrated that the molecular limb program in salamander limb progenitor cells is tuned to Shh-signaling as the primary input. Thus, Shh-signaling was sufficient to induce limb morphogenesis from forelimb progenitors in *Shh*-crispants. Interestingly, implantation of SHH-N beads (or FGF8 beads) could not induce ectopic limbs at stage 37-39 prior to when forelimb buds emerge from the flank. These experiments support that forelimb progenitors are specified much earlier during salamander development (*62*) and that these cells wait until endogenous Shh is activated which jumpstarts cell proliferation and activates downstream signaling. Future studies examining molecular players that may lie upstream of Shh (e.g., RA and Tbx5/4) will provide important evidence as to which signals actually establish the field and secondarily induce Shh expression in these cells. Classical experiments have shown that pieces of otic placode implanted into the flank can induce an ectopic limb (*70*) and the otic placode expresses Fgf8 among other factors. This raises the possibility that while salamander limb development may have shifted its reliance on Fgf-signaling for morphogenesis, it still may be a key factor for establishing the limb field.

Given the most recent phylogenetic models for relationships among extant tetrapods (*71, 72*), our findings support that urodeles possess a derived program for limb development relative to anurans and amniotes (Figure 6). This deviation appears to occur after establishment of limb field progenitors (*14*) and CRISPR/Cas9 knockout studies in newts show that upstream regulators of limb bud initiation like *Tbx5* are conserved among vertebrates (*73*). The results of this study in combination with our previous results showing movement of Fgf-signaling to the mesoderm (*3*) and work showing that the core long-range *Shh* enhancer (ZRS) is dispensable for limb development (*73*) supports that the molecular circuitry during limb outgrowth has been reconfigured in urodeles. Interestingly, a study in medaka (*Oryzias latipes*) showed that in addition to the canonical ZRS, a shadow ZRS (sZRS) controls *Shh* expression in fish (*74*) and only deletion of both enhancers resulted in the complete loss of *Shh* expression and loss of pectoral fins (*74*). Considered together, the results from these studies point towards the existence of a shadow enhancer for *Shh* in urodeles limbs similar to fishes. With the loss of a morphological and molecular AER (*3*), the limb development program in urodeles relocated into the mesoderm with a role for the ectoderm diminished (*75, 76*). The derived nature of the urodele limb program raises the intriguing hypothesis that the ability to regenerate an adult limb evolved in this lineage as opposed to limb regeneration being ancestral for tetrapods only to be lost in all the other major extant lineages (Figure 6). Functional studies using cyclopamine during normal limb regeneration in axolotls have shown contradictory phenotypes (*77, 78*). While an earlier study showed that Shh-signaling just controls anteroposterior patterning during limb regeneration (*77*), a later study using almost double the concentration of cyclopamine showed that Shh-signaling maintains *Fgf8* expression which is crucial for regenerative outgrowth (*78*). Moreover, the canonical AER-specific *Fgfs* (*Fgf8, 9* and *17*) are re-expressed in the limb mesenchyme during normal limb regeneration (*78*). In contrast to its relatively minor function during development, this study suggested that Fgf-signaling is required during limb regeneration where it functions in a positive feedback loop with Shh-signaling. Although many have argued on the basis of comparative gene expression data that an ancestral regeneration program exists for fins and limbs (*79-81*), these studies do not account for the fact that common gene expression profiles obscure functional relationships between signaling pathways. Regeneration of pectoral fins from endochondral amputations exist in some but not all Actinopterygian lineages. Moreover, regeneration of cartilaginous elements in adult lungfish fins resembles regeneration of cartilage spikes following amputation of *Xenopus* limbs. Although they are often ascribed limb regenerative ability as tadpoles, anurans can never regenerate a fully formed limb even as larvae (*82*) supporting that only urodeles, among extant amphibians, can regenerate an entire limb.

As such, our results underscore the need to expand limb development studies across a more diverse array of vertebrates, especially anamniotes. For instance, while the relatively few molecular studies in anurans support a mode of limb development more aligned with amniotes, the data also suggests an alternative dorsoventral patterning system may be in place (*52*). Additionally, although anurans appear to exhibit conserved ectodermal-mesodermal crosstalk (*83*), studies in the direct-developing frog *Eleutherodactylus coqui* indicate that a morphological AER is not required for compartmentalized ectodermal *Fgf8* expression (*84, 85*). Recent study that showed the absence of *Fgf8* during the development of bowfin pectoral fins with an AER further supports the plasticity in the limb/fin molecular program (*86*). Relative to amniote limbs which develop with input from the somites, amphibian limbs exhibit delayed development and a high degree of self-organization in that transplantation of limb buds to other parts of the body produce a relatively normal limb (*62*), an ability which may help support limb regeneration (*87*). Although other tetrapods exhibit high degrees of self-organization in the limb field, our results offer that shifting the limb program entirely into the mesenchyme and towards a more pronounced reliance on Shh-signaling to coordinate limb outgrowth may have permitted the self-organizing behavior of limb progenitors to execute patterning and ultimately, functional regeneration.

## Materials and Methods

### Animal husbandry and tissue harvest

Axolotls (*Ambystoma mexicanum*) (albino and wildtype) were acquired from our own laboratory colony. Chicken eggs (University of Kentucky, Department of Animal Sciences) were incubated to required stages. All procedures were conducted in accordance with, and approved by, the University of Kentucky Institutional Animal Care and Use Committee (IACUC Protocol: 2013–1174). For detailed methodology of animal husbandry and tissue harvest, refer Purushothaman et al., 2019.

Axolotl larvae were reared at 20–21 °C and larvae used for drug treatments, Alcian blue/Alizarin red staining, whole mount *in-situ* hybridizations, cell proliferation and cell death assays were anesthetized using 1x benzocaine (Sigma) and fixed overnight in 4% paraformaldehyde (PFA) at 4 °C. for qRT-PCR, larvae were anesthetized using 1x benzocaine, limb tissue samples were snap frozen and stored at -80°C until further use.

Chicken embryos (Single Comb White Leghorn) were incubated in 1502 Sportsman incubator, at 99.5 °F, with 40-50% humidity, harvested at HH30-HH32, fixed overnight in 4% PFA at 4°C and processed for various downstream assays.

### Drug treatments on axolotl larvae and chick embryos

Drug treatments on stage 39 axolotl larvae were done according to Purushothaman et al., 2019. Larvae were reared in 6 well plates and kept in dark throughout the experiment. A working stock of 5mg/ml of cyclopamine (Sigma) and 5 mg/ml of BMS-833923 (Cayman Chemical) was prepared in 100% ethanol and 0.6μl from this stock was added into 3ml 20% Holtfreter’s solution per well (1μg/ml final concentration). An equal amount of 100% ethanol (0.02%) was added into control wells. For earlier drug treatments, de-jellied axolotl embryos were treated with ethanol, cyclopamine or BMS-833923 at neural fold stage (stage 19/20) for 10 days. The solutions were replenished every two days and treatments lasted for 10 days.

Drug treatments on chicken embryos were done at HH14/15. 5ml of albumen was removed from the bunt end using a 5ml syringe and 22G1½needle. The eggs were windowed, and vitelline membrane around the limb bud was removed. The embryos were treated with 5μl of 1mg/ml solution of BMS-833923 in DMSO or DMSO (control) followed by 200μl of Ringers solution with 100U/ml Pen-strep. The window was covered with a scotch tape and the eggs were reincubated until harvest at HH30/32.

### Alcian blue and Alizarin red staining

Alcian blue and Alizarin red staining on axolotl larvae was done as previously described in Purushothaman et al., 2019. Fixed axolotl larvae were dehydrated in graded ethanol series and stained with 0.02% Alcian blue 8GX (Sigma Aldrich) in 70% ethanol and 30% glacial acetic acid for 3 hr to overnight. Larvae were rehydrated in graded ethanol series and then stained with 0.1% Alizarin Red (Sigma Aldrich) in 1%KOH overnight. Larvae were cleared in 1%KOH/glycerol series: 3KOH:1glycerol (imaged when cleared), 1KOH:1glycerol (1 day) and 1KOH:3glycerol (stored at room temperature).

Chicken embryos were harvested at HH30/32 and fixed in 100% ethanol for two days, stained with 0.1% Alcian blue 8GX (Sigma Aldrich) in 80% ethanol/20% acetic acid for one day and cleared in 1% KOH before imaging.

### Whole mount *in-situ* hybridization

Sense and antisense probes for *Ptch1, Grem1* and *Fgf8* axolotl genes were synthesized according to Purushothaman et al., 2019. Fixed axolotl larvae were dehydrated in graded methanol/PBT series stored in 100% methanol at -20°C until further use. Larvae were rehydrated in a graded methanol/PBT series and bleached with 6% H_2_O_2_/1x PBS for 1 hr under ice-cold conditions. Larvae were permeabilized with 20 μg/ml Proteinase K (Roche) in PBS for 7–10 min, fixed with 0.2% gluteraldehyde/4% PFA at room temperature and incubated overnight in hybridization buffer (5% Dextran sulphate, 2% blocking powder from Roche, 5X SSC, 0.1% TritonX, 0.1% CHAPS from Sigma Aldrich, 50% formamide, 1 mg/ml tRNA from Roche, 5 mM EDTA from Sigma and 50 μg/ml Heparin from Sigma) at 60 °C. The tubes were replaced with fresh hybridization buffer, 0.1–1 μg of probe was added into each vial and incubated at 60°C for two days. High stringency washes were done with 2X SSC/0.1% CHAPS thrice for 20 min each, 0.2X SSC/0.1% CHAPS 4 times for 25 mins each and with KTBT (15mM Tris-HCl pH 7.5, 150 mM NaCl, 10mM KCl and 1% Tween 20) twice for 5 min each. Larvae were blocked with 20% goat serum in KTBT for 3 hr, later treated with fresh blocking solution with an anti-Digoxigenin-AP, Fab fragment antibody (Roche) at 1:3000 dilution and incubated overnight at 4°C. Larvae/embryos were washed with KTBT 5 times for 1 hr each and then incubated in KTBT overnight at 4°C. Larvae/embryos were washed with NTMT (100mM Tris-HCl pH 9.5, 50mM MgCl_2_, 100mM NaCl and 1% Tween 20) and incubated in NBT/BCIP (Roche) solution (BCIP-0.17mg/ml, NBT-0.33mg/ml, 10% DMF in NTMT) till a signal developed with minimum background staining.

### qRT-PCR analysis

Stage 39 axolotl larvae were reared in 6-well plates in either 0.02% ethanol, 1μg/ml cyclopamine or 1μg/ml BMS-833923 till stage 46. Whole limbs were dissected from the body wall, immediately snap frozen and stored at -80 °C until RNA extraction. n=3 was used for each condition and each replicate represented a pool of limbs (both left and right) from 10 to 20 animals.

RNA was extracted using Trizol reagent (Invitrogen), cDNA was synthesized from 0.5 to 1μg RNA using SensiFast cDNA synthesis kit and qRT-PCR was performed using iTaq Universal SYBR Green Supermix (Biorad) (refer Purushothaman et al., 2019 for primer sequences). Melting curve was analyzed to confirm primer specificity.

*Rlp32* were used as the internal control/house-keeping genes for the experiments, respectively, since there was no significant fold change in the 2^-Ct^ values (*88*). 2^-ΔΔCt^ method was used to calculate the fold change values between control (ethanol) and treatment (cyclopamine or BMS-833923) groups (*88*).

### Guide RNA synthesis

Protocol for guide RNA synthesis was partially adapted from Fei et al., 2018. The mRNA coding sequence for *Shh* gene was accessed from https://www.axolotl-omics.org/search. DNA template oligos for guide RNA synthesis were designed using cloud-based informatics platform Benchling and oligos (20-mer or 18-mer) with high on-target and off-target scores were selected. Three DNA template oligos for separate *Shh*-guide RNAs were ordered from IDT with a 5’adapter and T7 promoter at the 5’ end and a 3’ overhang sequence complementary to the constant sequence at the 3’ end (refer Fei et al., 2018 for schematic diagram and Table S1 for sequences). The DNA template oligo for *Tyrosinase* (*Tyr*)-guide RNA ordered from with 5’ adapter, T7 promoter sequence, GG nucleotides at the 5’ end and a 3’ overhang sequence complementary to the constant sequence at the 3’ end (refer Fei et al., 2018 for schematic diagram and Table S1 for sequences). The DNA template oligo for *Tyr*-guide RNA was adapted from Fei et al, 2018.

The DNA template oligo was amplified using the Phusion DNA polymerase kit (NEB, cat# M0530S). Refer Table S1 for primer sequences. The reaction mix and PCR reaction were:

|  |  |
| --- | --- |
| Phusion GC buffer, 5x | 10µl |
| dNTPs (10mM) | 0.5µl |
| gRNA-fw2 (100µM) | 0.35µl |
| DR274-rev (100µM) | 0.35µl |
| gRNA oligo (100µM) | 0.25µl |
| Constant sequence (100µM) | 0.02µl |
| Phusion DNA polymerase (2U/µl) | 0.5µl |
| Nuclease free water | 38.03µl |
|  | 50µl |

### PCR Reaction

|  |  |  |
| --- | --- | --- |
| 98°C | 30sec | x 1 |
| 98°C | 20sec | x 35 |
| 60°C | 20sec |  |
| 72°C | 20sec |  |
| 72°C | 5min | x 1 |
| 4°C | forever | x 1 |

The PCR product was checked on a 1% agarose gel for single bands, purified (in 20μl water) using QIAquick PCR purification kit and quantified. The purified product was used for guide RNA synthesis (*in vitro* transcription) using Ambion MegaShortscript Kit T7; (cat#AM1354). The *in vitro* transcription step was as follows:

|  |  |
| --- | --- |
| T7 10X reaction buffer | 2μl |
| T7 ATP solution (75mM) | 2μl |
| T7 CTP solution (75mM) | 2μl |
| T7 GTP solution (75mM) | 2μl |
| T7 UTP solution (75mM) | 2μl |
| Template DNA from previous PCR (500-750ng) | xμl |
| T7 enzyme mix | 2μl |
| Water | make up to 20μl |

The reaction mix is incubated at 37°C overnight and guide RNAs were precipitated by phenol/chloroform method as follows: the reaction mix was transferred into a 2ml vial, 115μl water and 15μl Ammonium acetate stop solution (from Ambion MegaShortscript Kit T7) and 500 μl phenol+500μl chloroform mix were added, mixed till an emulsion formed, centrifuged at 13000 rpm for 1min, aqueous phase was transferred into a fresh 1.5ml vial, 2 volumes of ethanol was added and mixed well, chilled at -20°C for 15min, centrifuged at 13000rpm for 15min, supernatant was carefully discarded, tubes were allowed to dry under the hood and the RNA pellet was resuspended in ∼20μl water. The integrity of the guide RNAs was checked on a gel, quantified and stored in -70°C (as 2μl aliquots) until microinjections.

### Microinjections

Protocol for microinjection was partially adapted from Fei et al., 2018. The following injection mix was freshly prepared once the female started laying eggs:

|  |  |
| --- | --- |
| CAS9-NLS protein (PNA Bio, cat# CP03), reconstituted in 1x CAS9 buffer to 5mg/ml | 1μl |
| Guide RNA (4μg) | xμl |
| CAS9 buffer, 10x (200 mM HEPES, 1.5 M KCl, pH 7.5) | 0.9μl |
| Nuclease free water | make up to 10μl |

The glass capillary tubing (OD=1 mm, ID=0.58 mm, length=7.5 cm) was pulled (Sutter instruments Co., settings-heat:600, pull:50, vel:120, time-165) and the injection mix was loaded into it. The needle tip was calibrated using a stage micrometer so as to inject a volume of 5nl into each fertilized egg. For control injections (CRISPR control), injection mix minus guide RNA was injected into each single celled fertilized egg.

The single celled fertilized eggs were sterilized using 70% ethanol for 20 sec, rinsed and dejellied in 1x MMR/Pen-strep solution and transferred to 1x MMR/Pen-strep+20% Ficoll solution for microinjections. Post injection, the eggs were transferred into fresh 1x MMR/Pen-strep+20% Ficoll for 2hr and then transferred into 0.1x MMR/Pen-strep+5% Ficoll for 24hr at 18°C. The healthy embryos were transferred into 24 well plates with 0.1x MMR/Pen-strep solution and not disturbed for 7 days. Fresh 0.1x MMR+Pen-strep solution was added on the 8^th^ day and solution was replenished once in two days till final harvest.

### Genotyping

Protocol for genotyping was partially adapted from Fei et al., 2018. For genomic DNA extraction, 1mm tail clips from CRISPR control and guide RNA injected larvae was snap frozen in 1.5ml vials and stored at -80°C until further use. Later, into the 1.5ml vials 100μl 50mM NaOH was added, incubated at 95°C for 20min, 10μl 1M Tris pH 7.5 was added, mixed well, spun and quantified. The following PCR was performed to amplify the gene locus:

|  |  |
| --- | --- |
| Genomic DNA (200ng) | xµl |
| Taq reaction buffer, 10x | 2µl |
| dNTPs (10mM) | 0.3µl |
| Genotyping primer forward (10µM) | 1µl |
| Genotyping primer reverse (10µM) | 1µl |
| Taq DNA polymerase (5U/µl) | 0.1µl |
| Nuclease free water | make up to 20µl |

(Refer to Table S1 for genotyping primer sequences)

### PCR reaction

|  |  |  |
| --- | --- | --- |
| 94°C | 2min | x 1 |
| 94°C | 30sec | x 35 |
| 50 or 60 or 70°C | 30sec |  |
| 72°C | 30sec |  |
| 4°C | forever | x 1 |

The PCR product was checked on gel to verify single bands, quantified and sent out for Next Generation Sequencing (Amplicon-EZ, Genewiz) to sequence each gene locus.

### Bead experiments

Axolotl embryos were used at developmental stage 37 (*63*). Before any procedure, embryos were anesthetized using 1x benzocaine (Sigma) and placed on 1% agarose plates under a stereoscope microscope. For bead implantation, a tungsten wire was used to make an incision into the posterior ectoderm where the control and treated beads were slid into the presumptive trunk. The same procedure was used after three and six days to implant a second and third bead following the treatment. Axolotls were euthanized and fixed in 10% neutral buffered formalin (NBF, StatLab) for imaging.

Affi-beads were used (Affi-Gel Blue Media, cat#153-7301, 150-300um) with SHH-N protein (R&D, cat#1845-SH-025). Affi-beads were incubated in 1μl (0.25μg/μl or 0.5μg/μl) SHH-N and reconstituted in PBS with 0.1% BSA for 2hr at room temperature or overnight at 4°C. For FGF8 experiment, heparin/agarose beads (Sigma-Aldrich, H6508) were washed in PBS and then individually soaked in a 2μL drop of FGF8 protein solution for 2 hours at room temperature. Control beads were incubated in 1μl PBS with 0.1% BSA for 2hr at room temperature or overnight at 4°C. Prior to grafting, Fgf8-soaked beads were then transferred into a 2μL drop of 0.1% fast green dissolved in water in order to visualize the beads.

Rescue bead experiments were done on larvae injected with either *Shh*-guide RNA#3 or *Shh*-guide RNA#1, 2 and 3 with evident axolotl *Shh* cripant phenotypes like curved body axis, partial to complete cyclopia, no limb outgrowth etc. The beads were grafted when the CRISPR control larvae reached stage 44. Larvae were reared in 24-well plates in 0.1X MMR/Pen-strep solution. Larvae were anesthetized in 1x benzocaine for 1 min and transferred to 1% agarose plates with PBS with 0.1% BSA solution. A nick was made at the approximate limb field position using a tungsten needle and beads were grafted securely into the nicks. Protein-soaked bead was grafted into the right limb field and control bead was grafted into the left limb field. Larvae were placed back into the 24-well plates in fresh 0.1X MMR/Pen-strep solution. Beads were replaced once in 3-4 days and experimental larvae were harvested once CRISPR control larval limbs reached stage 46/48.

### Cell death and cell proliferation assays

Refer Purushothaman et al., 2019 for detailed protocols of cell proliferation and cell death assays in axolotls. Post-hatch larvae were reared in 6-well plates in 3 ml of either of the solutions: 0.02% ethanol, 1μg/ml cyclopamine or 1μg/ml BMS-833923 in 20% Holtfreters solution. CRISPR control and *Shh* crispants were reared in 6-well plates in 3 ml of 0.1x MMR/Pen-strep solution.

For cell proliferation assay, larvae were additionally treated with 0.1 mg/ml of EdU for 24hr when control larvae (ethanol treated or CRISPR control) reached stage 45, fixed overnight in 4% PFA, dehydrated in 1x PBS/methanol series and stored in 100% methanol at -20°C until further use. Larvae were rehydrated backwards through graded methanol series starting at 100% methanol and ending at 100% 1x PBS, treated with 2.5% trypsin (Gibco) for 10min, permeabilized with 20μg/ml of proteinase K in PBT for 7–10min, fixed in 100% acetone at -20°C for 10min, incubated in fresh click reaction solution (1x TRIS buffer saline, 4mM CuSO_4_ in 1x TRIS buffer saline, 2μl Alexa-flour-594 Azide (Life technologies), 1mM sodium ascorbate in 1x TRIS buffer saline) for 30min on a rocker in the dark, incubated in DAPI (1:1000 dilution) for 30mins, checked for fluorescence under a stereomicroscope and stored at 4°C in the dark until lightsheet imaging.

For cell death assay, larvae were transferred into 24-well plates and treated with 200μl of 5μM LysoTracker Red DND-99 (molecular probes) in Hanks BSS for 45min to 1hr at 20–21°C, fixed overnight in 4% PFA, dehydrated through a graded methanol/ Hanks BSS series and stored in 100% methanol at -20°C until imaging.

### Microscopy and image analysis

Whole mount images of limbs for Alcian blue/Alizarin red staining, limb size measurements, *in situ* hybridization, bead experiments, cell death assays, body axis analysis and eye position and pigmentation analysis were taken on an SZX10 light microscope (Olympus, Tokyo, Japan) using a DP73 CCD camera (Olympus). The microscope was equipped with CellSense software (CellSense V 1.12, Olympus corporation).

EdU stained stage 45 larval limbs were imaged using a Zeiss Lightsheet Z.1 (College of Arts and Science Imaging Centre, University of Kentucky). Refer Purushothaman et al., 2019 for detailed protocol of lightsheet microscopy for axolotl limb buds. Zen software (Zeiss) was used for imaging and samples were excited using 561 nm and 488 nm lasers. Arivis vision4D software (Arivis) was used for image processing. For total limb volume calculations, an object mask was hand drawn at each z-plane based on the DAPI signal to outline the limb. Red cell aggregate volume and total limb volume was calculated using previously standardized protocols and volume values in μm^3^ and voxel counts were given as outputs.

### Fiji analysis

Melanocyte pigmentation in the eyes of stage 46 tyrosinase crispants and control larvae was measured using Fiji software (NIH) after calibrations. Eye pigmentation was measured as pixel intensity and the ranges were as follows: pixel intensity=221 to 148 (high), =147 to 74 (moderate) and =73-0 (low).

### Statistics

All statistical analyses were performed using JMP (version Pro 12.10, SAS Institute Inc) and Microsoft Excel. Bar and Pie diagrams were made using Microsoft Excel.

For limb size between ethanol, cyclopamine and BMS-833923, one-way ANOVA followed by Tukey-Kramer HSD post hoc test was performed. Differences were considered significant if *P*<0.05.

For qRT-PCR data, the 2^-ΔΔCt^ method was used to calculate fold changes of genes between ethanol, cyclopamine and BMS-833923 groups. Calculations for mean Ct values, ΔCt values for treatment and control groups, ΔΔCt values between treatment and control groups, 2^-Ct^ and 2^-ΔΔCt^ fold change values were done using Microsoft Excel. One-way ANOVA followed by Tukey-Kramer HSD post hoc test was performed. Differences were considered significant if *P*<0.05.

For lightsheet data, red cell aggregate volume/limb volume (%) was calculated in Microsoft Excel (red cells=EdU positive cells). Post arcsin conversion, comparisons between control and treatment groups (ethanol vs. cyclopamine vs. BMS-833923 and CRISPR control vs. *Shh* crispant) were done by one-way ANOVA followed by Tukey-Kramer HSD post hoc test.

## Acknowledgments

We thank A. Cook, C. Hacker and J. Sarli for help with axolotl husbandry; J. R. Monaghan for training in CRISPR/Cas9 optimization; J. D. Currie for help with crispant analysis; E. M. Tanaka for access to previously unpublished axolotl sequences; M. Maden and all members of the Seifert lab for helpful discussions.

## Author contributions

Conceptualization: SP, AWS

Methodology: SP, BBL

Investigation: SP, BBL

Visualization: SP, BBL, AWS

Supervision: AWS

Writing—original draft: SP, AWS

Writing—review & editing: SP, AWS

## Data and materials availability

All data are available in the main text or the supplementary materials.

