## Supplemental Material for "*Sonic hedgehog* is essential for proximal-distal outgrowth of the limb bud in salamanders" (from another version of this preprint: version 1, 2021-09-16)

**This PDF file includes:**

Supplementary Text
Figs. S1 to S4
Tables S1 and S2

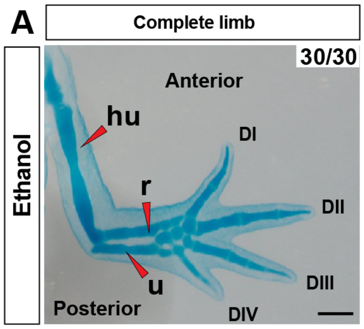


D

| Treatment | n= | Humerus | Radius/ulna | # Digits | # cases | % |
|---|---|---|---|---|---|---|
| Ethanol | 30 | Yes | Separate | 4 | 30/30 | 100 |
| Cyclopamine | 24 | Yes | None | 0 | 2/24 | 8 |
| | | Yes | ? | 0 | 1/24 | 4.2 |
| | | Yes | radius | 1 | 6/24 | 25 |
| | | Yes | radius | 1? | 4/24 | 16.7 |
| | | Yes | radius | 2 | 6/24 | 25 |
| | | Yes | radius | 2 | 3/24 | 12.5 |
| | | Yes | radius | 3 | 2/24 | 8.3 |
| BMS | 36 | Yes | radius | 1? | 1/36 | 2.8 |
| | | Yes | radius | 0 | 2/36 | 5.5 |
| | | Bud | None | 0 | 24/36 | 66.7 |
| | | No | None | 0 | 9/36 | 25 |

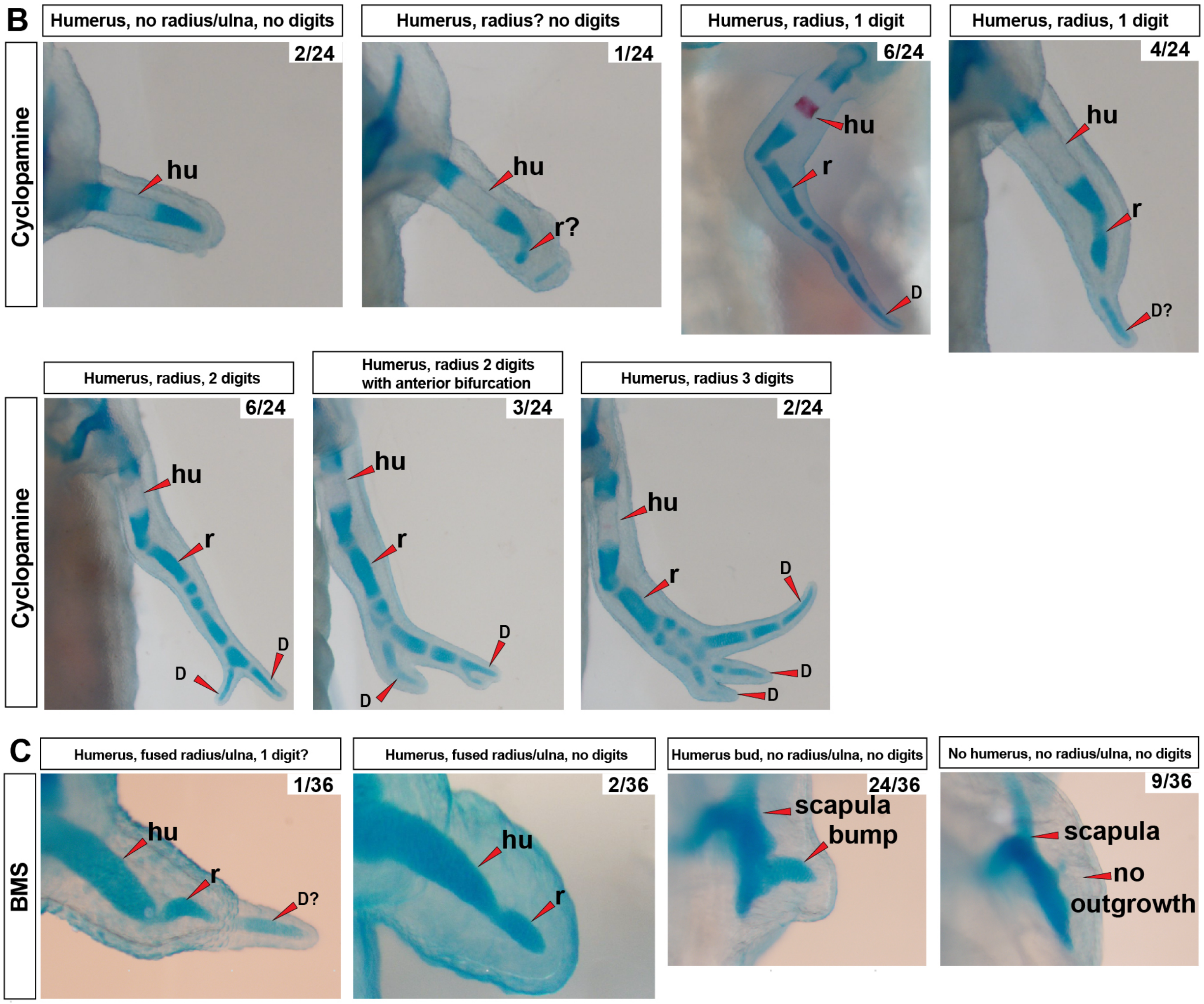


**Figure S1. Detailed range of limb phenotypes associated with ethanol, cyclopamine and BMS-833923 treatments. (A-C)** Alcian blue/Alizarin red staining for different limb phenotypes with case numbers in ethanol, cyclopamine and BMS treated larvae (limb n=30 for ethanol, =24 for cyclopamine and =36 for BMS). Scale bar=500 µm. **(D)** Table depicting treatments, limb phenotypes and percentage cases. hu, humerus; r, radius; ul, ulna and D, digits.

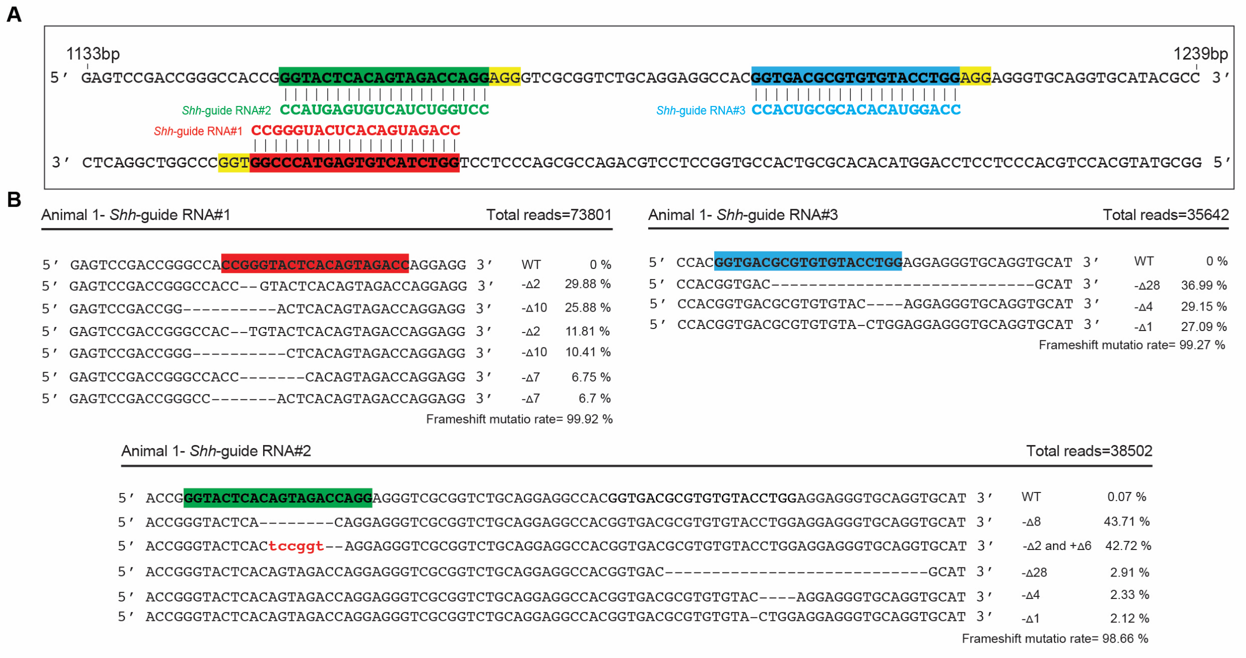

**Figure S2.** ***Shh* guide RNA gene targets and NGS results of *Shh* crispants.** **(A)** Target sequences for axolotl *Shh* (bold and highlighted in red, green and blue) and 20-mer *Shh*-guide RNAs #1 (red text), #2 (green text) and #3 (blue text) complementary to the target sequences. Protospacer Adjacent Motif or PAM (NGG) for each guide RNA target is highlighted in yellow. **(B)** Genotype of *Shh*-guide RNA injected larvae analyzed using AmpliconEZ with total number of reads, top 90% mutation types and frameshift mutation rates in the representative animals injected with either *Shh*-guide RNAs #1, #2 or #3. The regions highlighted in red (Animal 1-*Shh*-guide RNAs #1), green (Animal 1-*Shh*-guide RNAs#2) and blue (Animal 1-*Shh*-guide RNAs#3) are the target sequences for each guide RNA. Each animal had numerous mutation types: deletions are depicted as hyphens and insertions are in lower case and red text. WT: wild type sequence, -Δ: nucleotide deletions and %: percentage reads for each mutation type represented against it.

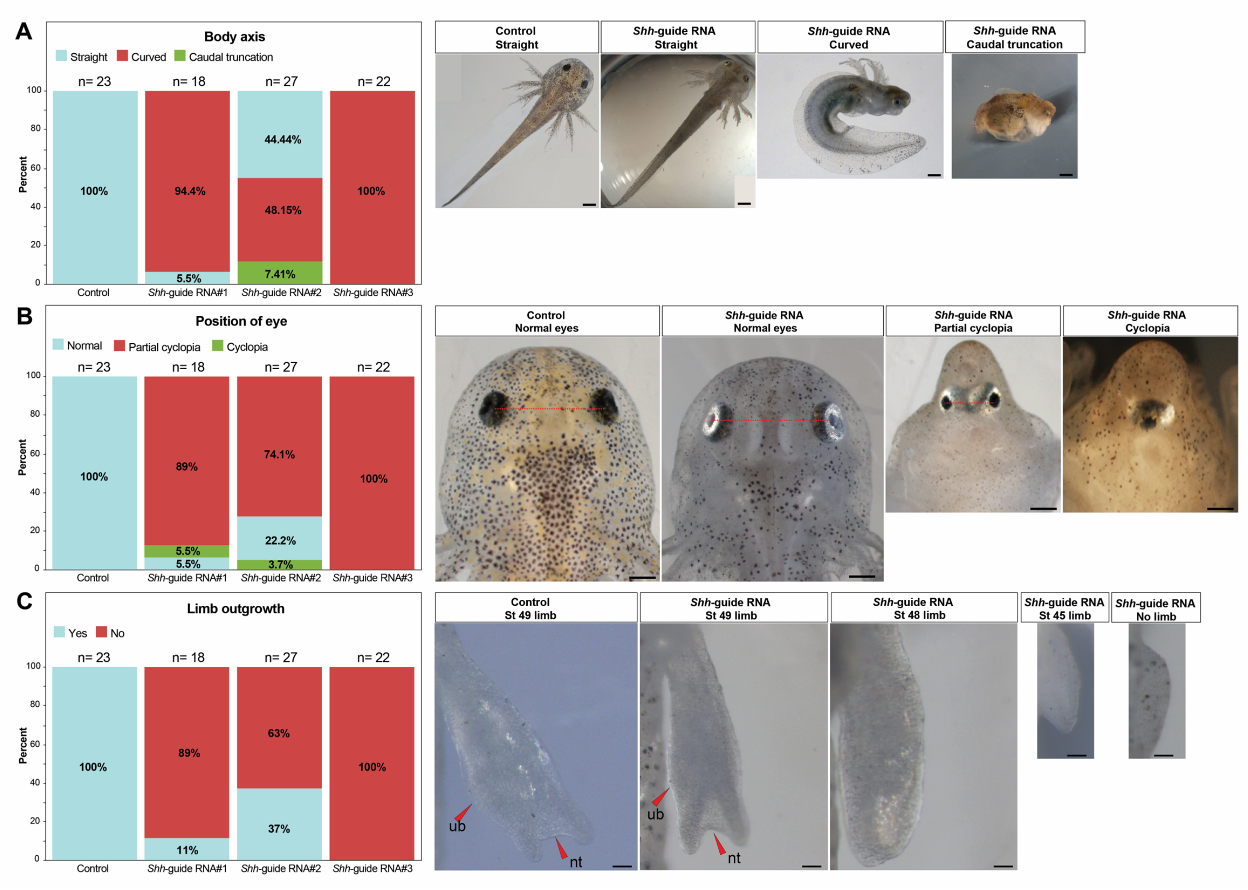


**Figure S3. Three separate *Shh*-guide RNAs result in highly overlapping phenotypes in F0 crispants.** **(A)** Various body axis phenotypes per guide RNA type in CRISPR control and *Shh* crispant larvae. **(B)** Various eye positions per guide RNA type in CRISPR control and *Shh* crispant larvae. Red dotted line: distance between the eyes. **(C)** Various limb phenotypes per guide RNA type in CRISPR control and *Shh* crispant larvae. n=23 for control larvae, n=18 for *Shh*-guide RNA#1 injected larva, n=27 for *Shh*-guide RNA#2 injected larvae and n=22 or *Shh*-guide RNA#3 injected larvae. Scale bar = 1 mm (for body axis), = 500 µm (for eye position) and =100 µm (limb stages). ub, ulnare bulge and nt, notch.

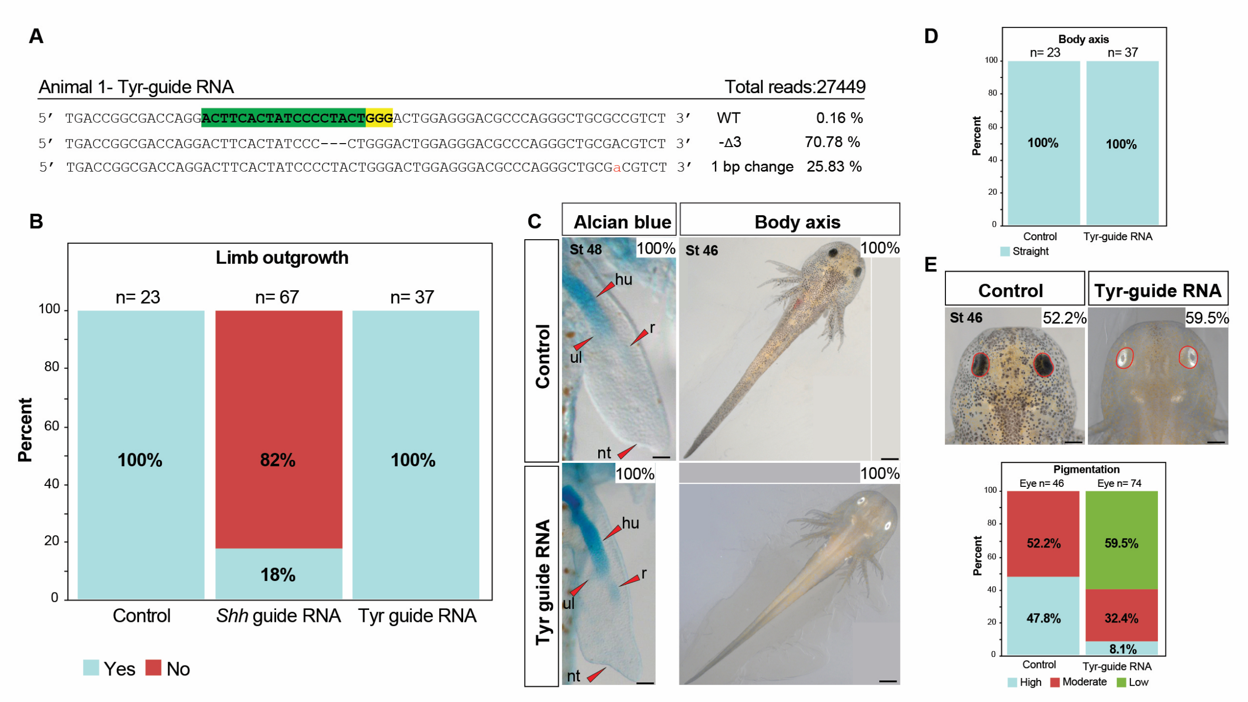


**Figure S5.** ***Tyrosinase* (*Tyr*) crispants do not show limb defects.** **(A)** Genotype of *Tyr* crispants analyzed using AmpliconEZ with total number of reads, top 90% mutation types and frameshift mutation rates in the representative animal. Deletions: hyphens and insertions: lower case and red text. WT stands for wild type sequence, -Δ represents nucleotide deletions and % reads for each mutation type is represented against it. **(B)** Percentage of CRISPR control, *Shh*- and *Tyr*-crispant larvae with limb bud outgrowth (n=23 for ethanol, =67 *Shh*-crispants and =37 *Tyr*-crispants). **(C, D)** Alcian blue staining (n=10 each) and body axis analysis (n=23 CRISPR control, =37 *Tyr*-crispants) for CRISPR control and *Tyr*-crispants. Scale bar = 1 mm (for body axis), = 500 µm (for Alcian blue staining). **(E)** Eye pigmentation (inside red dotted boarder) was calculated as pixel intensity using FIJI software and pixel intensity ranges were as follows: pixel intensity=221 to 148 (high), =147 to 74 (moderate) and =73-0 (low). n= 46 for CRISPR control and n=74 for *Tyr*-crispants. Scale bar = 500 µm. hu, humerus; ul, ulna; r, radius and nt, notch.

### Table S1. List of primers used in this study

| Experiment | Primers | Primer sequences 5' to 3' |
|---|---|---|
| WISH | Ptch1 Forward | GATCACAGAAGCTGGCTATGTAAAC |
| | Ptch1 Reverse | AAATATCAAGTCTGCAATCTTCGCG |
| | Fgf8 Forward | TTCACGTGCCTCACCTCCACTACCTCAGCA |
| | Fgf8 Reverse | ATATAAATTGTTCTTCTAAAAGTCCACTGG |
| | Gremlin1 Forward | TCGCCTGACACTACATAGC |
| | Gremlin1 Reverse | AGATGTGCAAAAGTTCAGAGAT |
| qRT-PCR | Fgf8 Forward | ATTAATTGTGGAAACGGACACCTTC |
| | Fgf8 Reverse | AATCAGCTTTCCCTTCTTGTTCATG |
| | Etv1 Forward | TCTTGGAAGAGTTCTTCTGAGTCAT |
| | Etv1 Reverse | CGTGTGAGAAATTGTAACGAGAGA |
| | Etv4 Forward | ACTATGCATACGATTCAGATGTTCC |
| | Etv4 Reverse | ATAGCCCTCCACGTTCATATACATT |
| | Gremlin1 Forward | GGACACCCAGAATACTGAGCA |
| | Gremlin1 Reverse | GTAGACCAATCGAAACATCCTGT |
| | Ptch1 Forward | TGTAGATCTGCTCCAATGCAAAC |
| | Ptch1 Reverse | CTGACCCGGAGTACTTGCAG |
| | Rlp32 Forward | AGGCTACTGGGAGTTTTAATAAGGA |
| | Rlp32 Reverse | AGATTACAGCACCCACTGTCTTTT |
| CRISPR/Cas9 | DNA template for Shh guide 1 | TTGAAATTAATACGACTCACTATAGGTCTACTGTGAGTACCCGGGTTTTAGAGCTAGAAATAGCAAGT |
| | DNA template for Shh guide 2 | TTGAAATTAATACGACTCACTATAGGTACTCACAGTAGACCAGGGTTTTAGAGCTAGAAATAGCAAGT |
| | DNA template for Shh guide 3 | TTGAAATTAATACGACTCACTATAGGTGACGCGTGTGTACCTGGGTTTTAGAGCTAGAAATAGCAAGT |
| | DNA template for Tyr guide | TTGAAATTAATACGACTCACTATAGGACTTCACTATCCCCTACTGTTTTAGAGCTAGAAATAGCAAGT |
| | Constant sequence | AAAAGCACCGACTCGGTGCCACTTTTTCAAGTTGATAACGGACTAGCCTTATTTTAACTTGCTATTTCTAGCTCTAAAAC |
| | gRNA-fw2 primer | TTGAAATTAATACGACTCACTATAGG |
| | DR274-rev | AAAAGCACCGACTCGGTGCCAC |
| | Shh genotyping primer forward | CGCGGCCCACCTCCTCTTTG |
| | Shh genotyping primer reverse | CGCCCAGCTGTGCTCCTCTATG |
| | Tyr genotyping primer forward | GCTCTTCCTGCTGCTCTGGGAG |
| | Tyr genotyping primer reverse | CTGCCAAGAAGCGAAGAAGGAG |

#### Table S2. NGS results

| Animal Sample | Target locus | Target Reads | Mutant Reads | Mutant % | Genotype | Frameshift Mutant Reads | Frameshift Mutant % |
|---|---|---|---|---|---|---|---|
| Control | *Shh* | 73144 | 917 | 1.25 | Homozygous WT | 129 | 0.18 |
| *Shh*-guide RNA#1 | *Shh* | 73801 | 73801 | 100 | Homozygous Mutant | 73739 | 99.92 |
| *Shh*-guide RNA#2 | *Shh* | 38502 | 38467 | 99.91 | Homozygous Mutant | 37987 | 98.66 |
| *Shh*-guide RNA#3 | *Shh* | 35642 | 35642 | 100 | Homozygous Mutant | 35382 | 99.27 |
| Control | *Tyrosinase* | 38044 | 57 | 0.15 | Homozygous WT | 53 | 0.14 |
| *Tyr*-guide RNA#1 | *Tyrosinase* | 27449 | 19584 | 71.35 | Heterozygous | 34 | 0.12 |
